## Supplemental Figure S1 and supporting text for "The Kv2.2 channel mediates the inhibition of Prostaglandin E2 on glucose-stimulated insulin secretion in pancreatic β-cells"

**This file includes:**

**Supplemental Figure S1. Generation of Kv2.2 knockout mice.**

**Supplemental Figure S2. Validation of EP receptor antibody specificity in INS-1(832/13) cells using siRNA knockdown.**

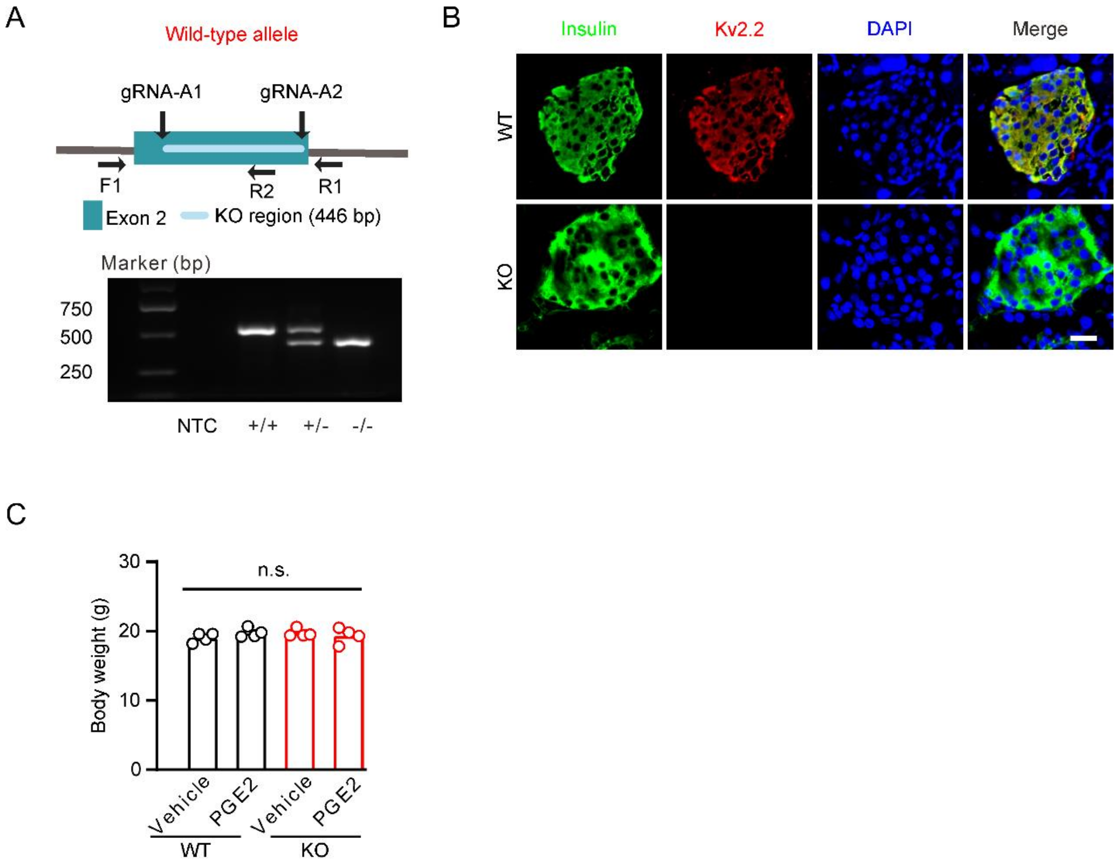


**Supplemental Figure S1. Generation of Kv2.2 knockout mice.**

**(A)** Top, cartoon of the Kv2.2 targeted gene disruption. Bottom, PCR products generated from wild-type (+/+, 521 bp), heterozygous (+/−, 521 bp and 415 bp), and homozygous knockout (−/−, 415 bp) mice with primers (F1 and R1) specific to the surrounding Kv2.2 gene disruption sequence and one primer (R2) specific to the targeting sequence. NTC, no template control. **(B)** Representative immunofluorescence images showing expression of Kv2.2 channels in wild-type and Kv2.2$^{-/-}$ knockout mouse pancreatic islet β-cells. Scale bar, 20 μm. **(C)** Kv2.2 knockout did not alter the body weight of animals.

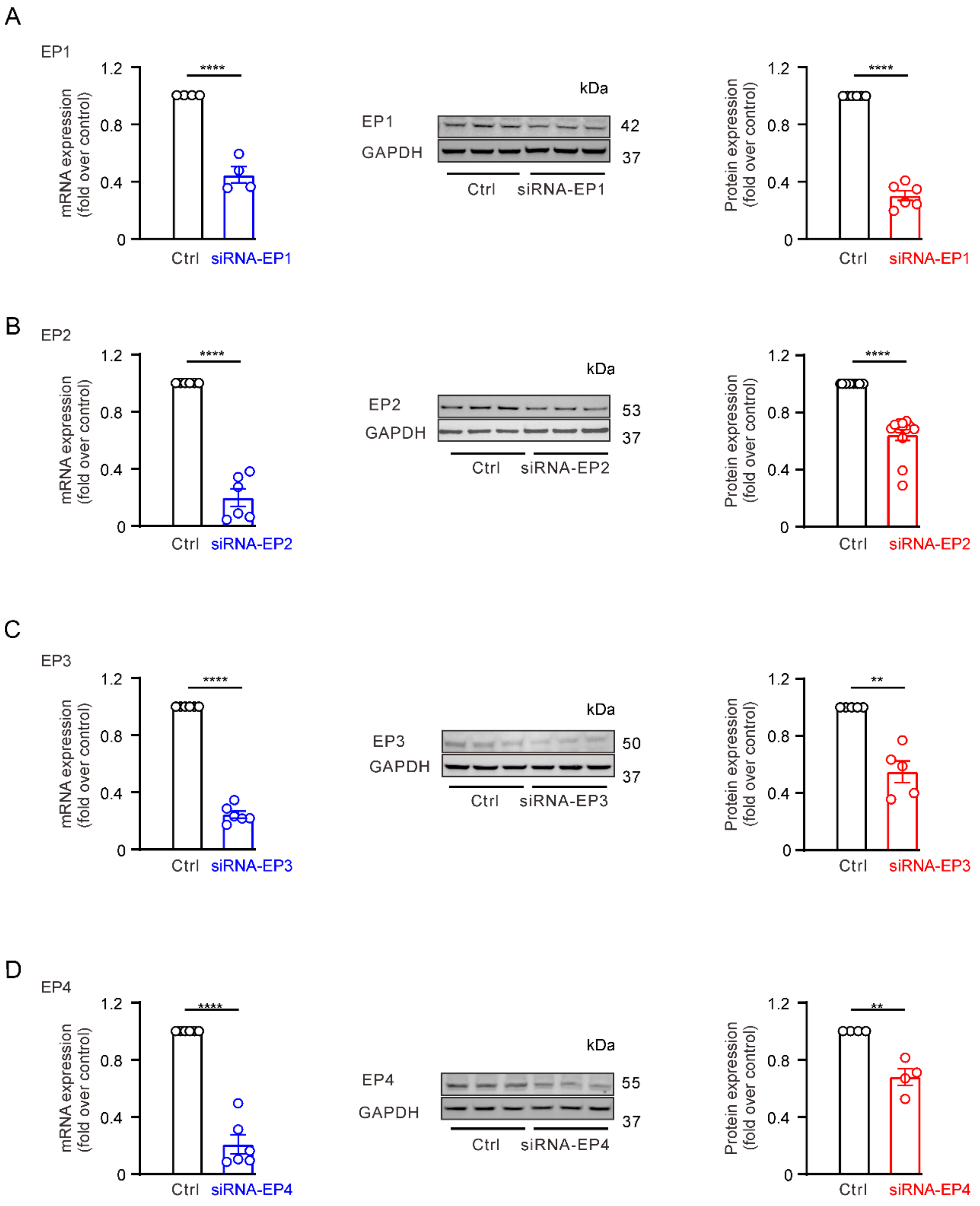


**Supplemental Figure S2. Validation of EP receptor antibody specificity in INS-1(832/13) cells using siRNA knockdown.**

**(A)** Left: Quantification of mRNA expression following siRNA targeting of EP1 receptors (siRNA-EP1). Middle: Representative western blot images showing the effect of siRNA-EP1 on EP1 protein levels. Right: Quantification of EP1 protein expression after siRNA-EP1 treatment. mRNA expression: n = 4, ****p < 0.0001; protein expression: n = 6, ****p < 0.0001; Two-tailed *t*-test.

**(B)** Similar to panel A, but assessing the effect of siRNA targeting EP2 (siRNA-EP2) on EP2 receptor expression. mRNA expression: n = 6, ****p < 0.0001; protein expression: n = 13, ****p < 0.0001; Two-tailed *t*-test.

**(C)** Similar to panel A, but assessing the effect of siRNA targeting EP3 (siRNA-EP3) on EP2 receptor expression. mRNA expression: n = 6, ****p < 0.0001; protein expression: n = 5, **p = 0.0042; Two-tailed *t*-test.

**(D)** Similar to panel A, but assessing the effect of siRNA targeting EP4 (siRNA-EP4) on EP2 receptor expression. mRNA expression: n = 6, ****p < 0.0001; protein expression: n = 4, **p = 0.0017; Two-tailed *t*-test.

The siRNA sequences targeting the four EP receptors are: siRNA-EP1 (forward, reverse):5′-GGUCACUACGAGCUACAGUTT and 5′-ACUGUAGCUCGUAGUGACCTT; siRNA-EP2(forward, reverse): 5′-GUCUGCGUCAUCCAUCACUTT and 5′- AGUGAUGGAUGACGCAGACTT; siRNA-EP3(forward, reverse): 5′- CUGGGUGCUGUGUCCAACGTT and 5′-CGUUGGACACAGCACCCAGTT; siRNA-EP4(forward, reverse): 5′-GGGCCUGUCAUUUCCUGGGTT and 5′- CCCAGGAAAUGACAGGCCCTT. All sequences were purchased from GenePharma (Suzhou, China). siRNAs were transfected using Lipofectamine™ 3000. Cells were collected at 48 hours and 72 hours post-transfection for quantitative real-time polymerase chain reaction (qRT-PCR) and Western blot experiments to validate knockdown efficiency. The “n” value represents the number of independent biological replicates.
